# Unified Analysis of Global and Focal Aspects of Absence Epilepsy via Neural Field Theory of the Corticothalamic System

**DOI:** 10.1101/339366

**Authors:** Dong-Ping Yang, P. A. Robinson

## Abstract

A physiology-based corticothalamic model is investigated with focal spatial heterogeneity, to unify global and focal aspects of absence epilepsy. Numerical and analytical calculations are employed to investigate the emergent spatiotemporal dynamics induced by focal activity as well as their underlying dynamical mechanisms. The spatiotemporal dynamics can be categorized into three scenarios: suppression, localization, and generalization of the focal activity, as summarized from a phase diagram vs. focal width and characteristic axon range. The corresponding temporal frequencies and spatial extents of wave activity during seizure generalization and localization agree well with experimental observations of global and focal aspects of absence epilepsy, respectively. The emergent seizure localization provide a biophysical explanation of the temporally higher frequency but spatially more localized cortical waves observed in genetic rat models that display characteristics of human absence epilepsy. Predictions are also presented for further experimental test.

**Author Summary:** Absence epilepsy is characterized by a sudden paroxysmal loss of consciousness accompanied by oscillatory activity propagating over many brain areas. Although primary generalized absence seizures are supported by the global corticothalamic system, converging experimental evidence supports a focal theory of absence epilepsy. Here we propose a dynamical mechanism to unify the global and focal aspects of absence epilepsy, with focal absence seizures associated with seizure localization, and the global ones associated with seizure generalization. Our corticothalamic model is used to investigate how seizure rhythms and spatial extents are related in these two different aspects of absence epilepsy. The results account for the difference of the experimentally observed seizure rhythms and spatial extents between humans and genetic rat models, which has previously been used to argue against the validity of such rats as animal models of absence epilepsy in humans.

## Introduction

Absence epilepsy is an idiopathic nonconvulsive generalized epilepsy, that displays a sudden paroxysmal loss of consciousness accompanied by abnormal brain oscillatory activity: 2.5 — 4 Hz ‘spike-and-wave’ discharges (SWDs) in the electroencephalogram (EEG), electrocorticogram (ECoG), and local field potentials (LFPs) [1–3]. These oscillations propagate rapidly over many brain areas and reflect the macroscopic dynamical properties of neuronal populations. Many experimental results led to the corticothalamic theory that interactions between cortex and thalamus generate absence seizures [3–8]. However, converging evidence of animal ‘models’ that display SWDs similar to those of absence epilepsy in humans [9–13] has shown that absence seizures can be triggered from a specific cortical focus, leading to a focal theory of absence epilepsy [2]. Also, an accurate source analysis from dense-array surface electrodes suggested that absence seizures in human patients were not truly ‘generalized’, with immediate and global cortical involvement, but rather were initiated in specific cortical regions, then propagated to the whole cortex within milliseconds [14, 15]. It remains a hot debate between the global and focal aspects of absence seizures [8,16–20].

Two genetic strains of rats: the Wistar Albino Glaxo/Rijswijk (WAG/Rij) and Genetic Absence Epilepsy Rats from Strasbourg (GAERS), have been well estabilshed as animal ‘models’ of absence epilepsy in humans, displaying similarities of simultaneous electro-clinical signs and performance prediction based on pharmacological data [21]. However, the frequency of SWDs is different: 7 — 11 Hz in rats [9, 11] and 2.5 — 4 Hz in cats, rhesus monkeys, or humans [1, 8, 17]. Although there is no *a priori* reason why the SWD frequency must be the same in all species, to our best knowledge, it is still unknown how a spontaneous 7 — 11 Hz rhythm can be initiated from and sustained in a cortical focus, and then spreading over other cortical regions via the so-called somatosensory-thalamo-cortical network and cortical propagation [3, 9, 11, 17]. Actually, the cortex and the thalamus form one complex oscillatory network, providing a resonant circuit to amplify and sustain the SWDs, with the resonance considered as an emergent property of the corticothalamic system [3, 22]. So the higher SWD frequency in rats may be accounted for by smaller brain sizes and shorter axons [23].

However, *spatial extent* is another significant difference of absence seizures between rats and humans. In rats, higher frequency SWDs were generated with a cortical ‘focus’ found within the perioral subregion of the somatosensory cortex [9]. During spontaneous absence seizures, the cortical focus drives widespread corticothalamic networks, which are responsible for the immediate onset, rapid propagation and high synchrony of SWDs. Specifically, the simultaneous electrophysiological recordings (LFPs, ECoGs, or EEG with simultaneous fMRI) in WAG/Rij rats have shown that SWDs are intense in the cortical focus, but spare in the occipital cortex, and there was a decreasing amplitude along the propagation direction on the cortex [2, 9, 24]. Thus, SWDs in WAG/Rij rats are more spatially localized than seizure activities in humans, which are more generalized over the whole cortex [8, 14]. It is also commonly found in experiments that lower-frequency resonances are much more generalized in space, than higher-frequency oscillations which are more localized to restricted cortical areas [25, 26]. So it is essential to investigate whether and how the spatial extents of seizure activities relate to the observed oscillations.

To the end, this paper explores the conditions for focal activity to be suppressed, to remain localized, or to generalize and spread over the whole brain. We theoretically investigate the spatiotemporal dynamics of the coupled cortex and thalamus using a physiology-based corticothalamic model with focal spatial heterogeneity, exploring the role of cortical propagation. We first introduce the detailed model, and numerical simulations and analytical methods. Then we present the phase diagram of the spatiotemporal dynamics vs. focus width and characteristic axon range. Specifically, the focal activity with a relative small width can be suppressed, while the one with a relative large width can generalize over the whole system with ~ 3 Hz traveling waves, and the one with a relative moderate width limits in the focal region without generalizing over normal region, but with a ~ 10 Hz rhythm. Here, in the focal aspect, absence seizures represented by electro-clinical symptoms are associated with seizure localization, while in the global aspect, absence seizures are associated with seizure generalization. Their spatiotemporal properties such as spatial extents and temporal frequencies are comparable with experimental observations in humans and rats. The resulting cortical waves have robust frequencies, and their underlying dynamical mechanisms are discovered by employing eigenvalue spectra and corresponding eigenmodes at critical states. Thus we uncover a unified dynamical mechanism for the global and focal aspects of absence epilepsy. We conclude with experimentally testable predictions and further possible extensions.

## Materials and Methods

Our corticothalamic model is described, following previous work [27], and specified here with focal spatial heterogeneity. Then, we describe the simulation methods, the analytical calculations of steady states and their linear stability analysis.

### Corticothalamic model

The corticothalamic system can be described by a continuum approach at the macroscopic level. Large-scale neural activities are determined by interactions between several neural populations, notably excitatory and inhibitory cortical neurons and thalamus, including reticular and relay nuclei. A schematic of these populations is presented in Fig. 1(A), with excitatory, inhibitory, reticular, and relay neurons represented by e, *i, r, s,* respectively. In such a model [27], the variables represent local mean values at position *r* = (*r_x_,r_y_*) in a 2D space: the local mean cell-body potential *V_a_*, the mean firing rate *Q_a_*, and the propagating axonal field *ϕ_a_*, for *a* = *e,i,r,s.* The physical interrelationship of these collective state variables is summarized in Fig. 1(B). The space can be scaled by the system length L to unify the human (*L* = 0.5 m) and rat (*L* = 0.025 m) brains into a unique framework here, with assumption of linear proportionality of other spatial properties in these two systems.

**Fig 1.**
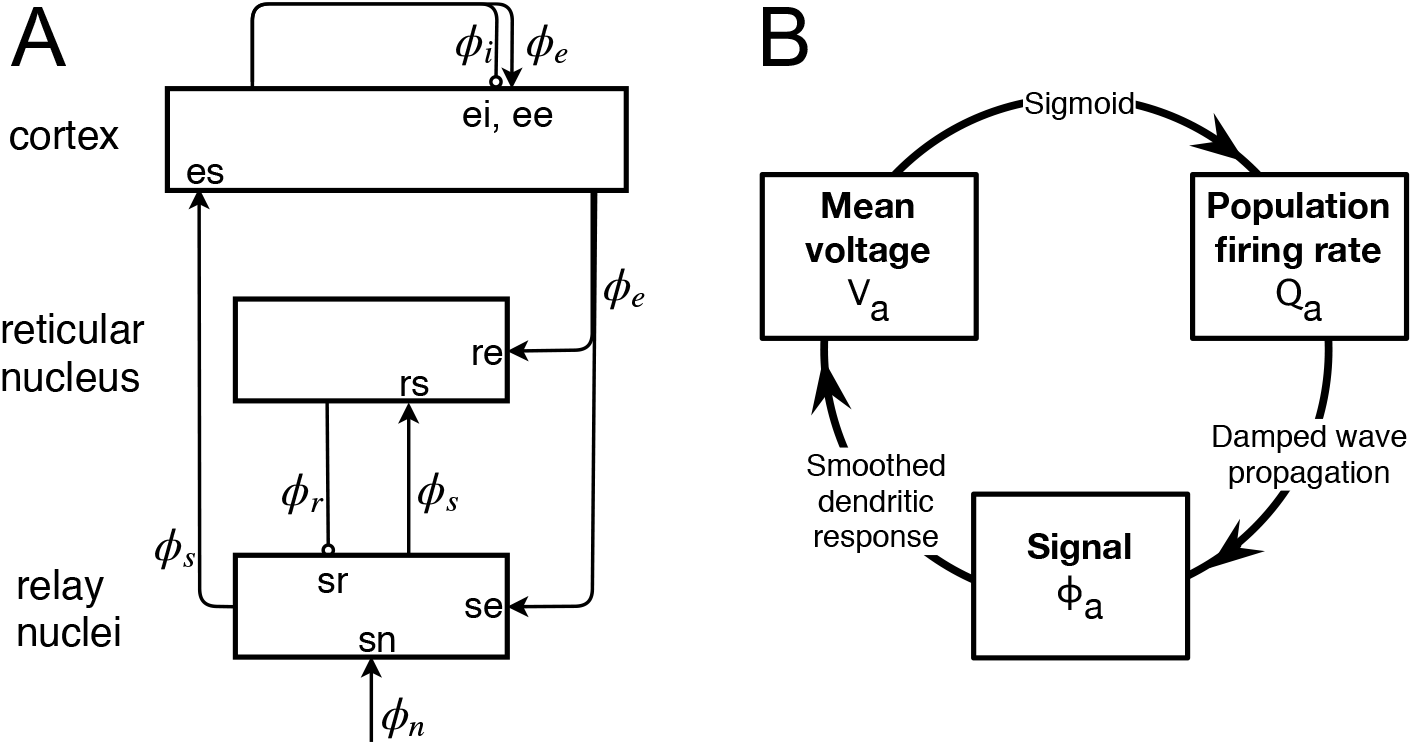
The corticothalamic model. (A) Schematic of the corticothalamic interactions, showing the locations *ab* at which couplings act. (B) The physical interrelationship of the system variables: *V_a_, Q_a_*, and *ϕ_a_*.

Firstly, the firing rate *Q_a_* is a sigmoid function of the potential *V_a_*, with

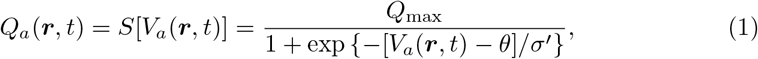

where *Q*_max_ is the maximal firing rate, *θ* is the mean firing threshold, and 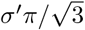 is the standard deviation of the difference between the steady state *V_a_* and the threshold *θ*.

Secondly, neuronal firing generates a field signal *ϕ_a_* and sends it through the extended axons toward other populations approximately according to the damped wave equation [28, 29]

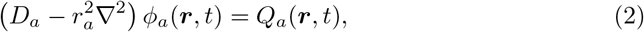

with

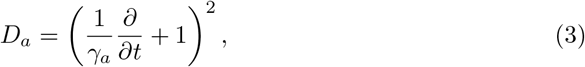

where *γ_a_* = *υ_a_/r_a_* governs the damping of propagating waves, and *r_a_* and *v_a_* are the characteristic range and conduction velocity of the axons of population *a*, respectively [29].

Finally, each population’s potential *V_a_* results when synaptic inputs from various types of afferent neurons are summed after being filtered and smeared out in time due to synaptic neurotransmitter, receptor dynamics, passage through the dendritic tree and the effects of soma capacitance. So

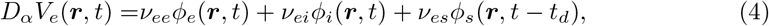

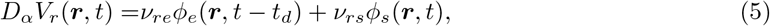

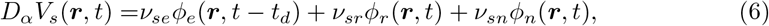

with the synaptodendritic operator

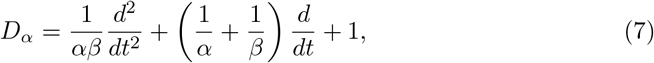

where *α* and *β* are the mean decay and rise rates of the soma response to an impulse arriving at a dendritic synapse [29].

Input from the thalamus to the cortex and feedback from the cortex to the thalamus are delayed by a propagation time *t_d_*. In Eqs (4) —(6), *ν_ab_* is the synaptic connection strength to population *a* from population b, and *ϕ_n_* is the ascending input from brainstem, which can be arbitrary external signals or approximated as white noise, but here is set to a constant for calculating phase diagrams and bifurcation diagrams. Figure 1(A) also presents the excitatory-inhibitory cortical loop, thalamic loop and corticothalamic loop in the model.

All populations except the excitatory population have very short axons, which lets us set *r_a_* ≈ 0 and *γ_α_* ≈ ∞ in (3), yielding *ϕ_α_* = *Q_a_* for *a* = *i,r, s* [27]. Besides, intracortical connectivities are found to be proportional to the numbers of synapses involved [30], so one finds *V_e_* = *V_i_* and *Q_e_* = *Q_i_* [27, 31], which allows us to concentrate on excitatory quantities, while implicitly retaining inhibitory effects on the dynamics. Thus the model includes 16 parameters: *Q*_max_, *θ*, *σ′*, *α, β, γ_e_*, *t_d_, r_e_, ν_ee_, ν_ei_, ν_es_, ν_se_, ν_sr_, ν_sn_ϕ_n_, ν_re_, ν_rs_*, which are enough to allow realistic representation of the most salient anatomy and physiology, but few enough to yield useful interpretations.

### Model of focal spatial heterogeneity

In WAG/Rij rats with absence seizures, the excitatory dendrites in the focal region have larger total dendrite length, larger mean length of a dendritic segment, and larger size of dendritic arbor, than those outside epileptic area [17, 21]. Additionally, the excitatory-inhibitory ratio of neuron numbers in focal cortical areas is larger than that in other areas and the efficiency of GABA-ergic inhibition is impaired [21]. Lesion studies also demonstrate that an excitable region is not sufficient for the occurrence of SWDs and indicate that some thalamic nuclei seem to be important for SWD occurrence [7], as borne out by theory and simulations [27, 32]. So all the above suggest that we use the corticothalamic model with focal spatial heterogeneity, which allows us to investigate the effect of cortical propagation on the spatiotemporal dynamics, and to study how the initially focal activity can be suppressed, remains localized, or propagate over the whole brain to produce secondary generalized absence seizures.

The above anatomical facts [17, 21] imply that a higher value of *ν_se_* in the focal area is suitable to describe the cortical focal activity, with *ν_se_* peaking at the center and smoothly decreasing to the edge, as shown in Fig. 2(A); *ν_se_* describes the excitatory influence of cortical pyramidal cells on the specific thalamic nuclei. The choice of *ν_se_* is also supported by prior implications of excitatory corticothalamic feedback in the pathophysiology of generalized absence seizures [9, 24, 34, 35]. For simplicity, the spatial profile is set as an isotropically symmetric Gaussian function

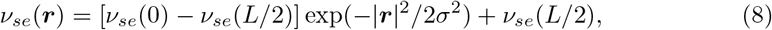

with *σ* characterizing the pathological width, *ν_se_*(*L*/2) the background normal value and *ν_se_*(0) the pathological value. Figure 2(B) shows profiles along *r_x_/L* at *r_y_* =0 with various *σ/L*. All other parameters are spatially invariant and given in Table 1.

**Fig 2.**
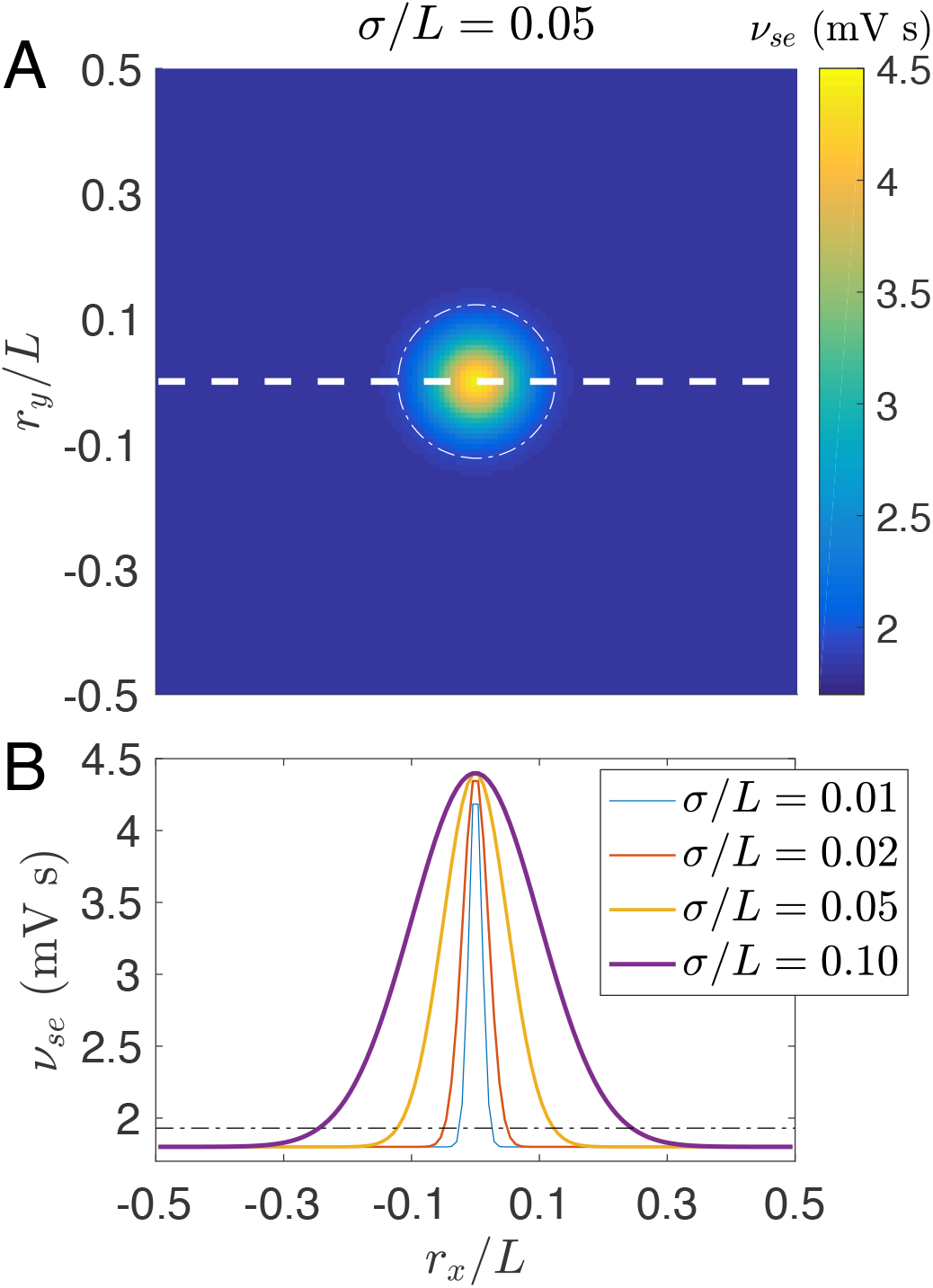
(Color online) Spatially heterogeneous corticothalamic model. (A) Schematic of the 2D spatial model with periodic boundary conditions; Each position *r* = (*r_x_,r_y_*) represents a corticothalamic loop with *ν_se_(r)* determined by a spatially Gaussian profile given in Eq. (8) with *σ/L* = 0.05, *ν_se_*(*L*/2) = 1.8 mV s and *ν_se_*(0) = 4.4 mV s. The white dashed line indicates the position of the spatial profiles in (B). (B) Spatially Gaussian profiles *ν_se_(r)* along *r_x_* at *r_y_* = 0 with various *σ/L*, as indicated in the legend. The white dash-dotted circle in (A) and the black dash-dotted line in (B) indicate the threshold 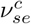 for transition to absence seizure in the spatially homogeneous case. They are also used to define the focal area inside and the normal region outside.

**Table 1.** Nominal parameter values in spatially homogeneous model from [32, 33].

| Parameter | value | Unit | Parameter | value | Unit |
| --- | --- | --- | --- | --- | --- |
| $Q_{\max}$ | 250 | $s^{-1}$ | $\nu_{ee}$ | 1.0 | mV s |
| $\theta$ | 15 | mV | $-\nu_{ei}$ | 1.8 | mV s |
| $\sigma'$ | 6 | mV | $\nu_{es}$ | 3.2 | mV s |
| $\alpha$ | 50 | $s^{-1}$ | $-\nu_{sr}$ | 0.8 | mV s |
| $\beta$ | 200 | $s^{-1}$ | $\nu_{sn}\phi_n$ | 2.0 | mV s |
| $\gamma_e$ | 100 | $s^{-1}$ | $\nu_{re}$ | 1.6 | mV s |
| $t_d$ | 40 | ms | $\nu_{rs}$ | 0.6 | mV s |

The isotropic symmetry is limited by boundary, which has little effect on the symmetry if we choose *σ/L* ≪ 1 and *r_e_/L* ≪ 1 but still brings edge effects on the spatiotemporal dynamics, as we will demonstrate in the results.

### Simulation methods

The model is simulated using the recently published NFTsim code written in C++ [36] to investigate the spatiotemporal dynamics [27, 32, 33], which solves our damped and retarded 2D wave equation in Eqs (2) and (4)–(6) with given initial conditions and boundary conditions.

In the numerical simulation, the 2D space (*r_x_,r_y_*) is divided into a 120 × 120 grid with *L* = 0.5 m for humans and *L* = 0.025 m for rats, and grid point spacing *δr_x_/L* = *δr_y_/L* = 1/120. We choose periodic boundary conditions and an initial condition that each spatial point is assigned with a *τ_d_* length time series of a random constant.

In NFTsim, numerical integration is performed using a fourth-order Runge-Kutta integrator. A cubic-spline interpolator is employed in order to estimate the time-delayed values of the midpoints required for the Runge-Kutta algorithm. A small time step (δt = 0.1 ms) is chosen to satisfy the Courant condition. Long (~ 1000 s) simulations are performed to guarantee the system reaching its stationary state.

Noise is ignored in the simulations for phase diagrams and bifurcation diagrams, which simplifies our analysis so as to focus on the pattern formation and their underlying dynamical mechanisms. We have checked that a small noise has no significant effect on our findings.

### Steady states and linear stability analysis

To get more insight of the underlying mechanisms for various spatiotemporal dynamics, we are introducing here analytical methods how to derive the steady states, their linear stability and eigenmodes.

As introduced above, the system has a radial symmetry with *σ/L* ≪ 1 and *r_e_/L* ≪ 1, leading to the replacement of *r* by *r* = |*r*| to simplify our analysis. Then the spatial interaction in radial coordinates can be rewritten as

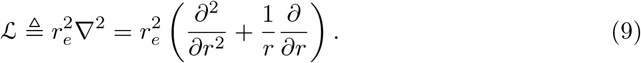

As proposed in Ref. [37], the steady states can be obtained by integrating Eqs (2), and (4) —(6) inward towards *r* = 0, starting at a large *r*, where the system can be linearized with r to be

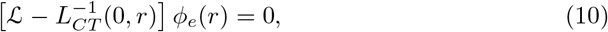

with the function 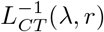 given by Eq. (34) in Appendix A, which describes the resonance 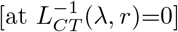 of the corticothalamic loop at r and each eigenmode with eigenvalue λ. The stationary state converges asymptotically to the solution at large *r*, and the appropriate boundary condition at *r* = 0 is

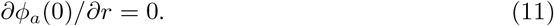

Thus one of its general solutions, converging at 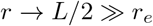, is

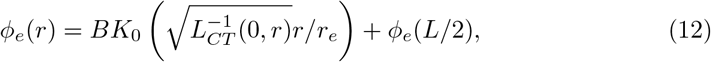

where *K*_0_ is a modified Bessel function of the second kind and *B* is an undetermined real constant. The choice of B uniquely parametrizes the stationary solution to the nonlinear equations (2), and (4) –(6). So our task is to search for the value of *B* for which the solution 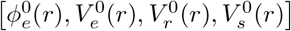 obeys the boundary condition (11).

To analyze linear stability of the steady state, a small dimensionless perturbation can be introduced as follows:

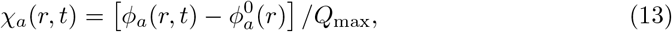

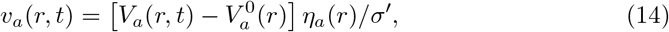

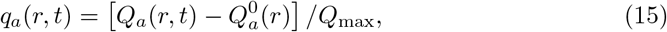

with 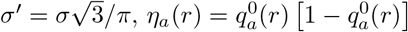, and 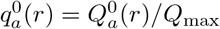 for *a* = *e, r, s*. The perturbations can be expanded in sums of eigenmodes as follows:

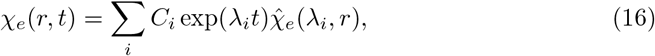

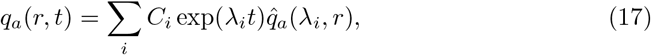

each of which obeys

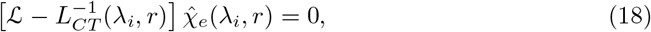

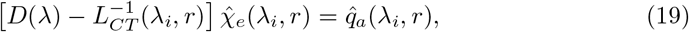

with the function *D*(λ) given by Eq. (28) in the Appendix A as well. Again, a unique bounded solution for 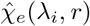 at *r* − *L*/2 is

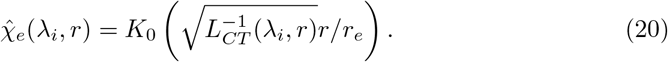

Note that there is no constant *B* as in the steady state equation (12) because it has been absorbed into *C_i_*. So now our task is to search for the eigenvalue *λ_i_* as well as the corresponding perturbation eigenmode. The largest Re(λ_i_) determines the linear stability of the steady state.

## Results

Our results are presented starting from the spatially homogeneous case. Then the corticothalamic system with initial focal activity is investigated by numerical simulations to study how cortical propagation can suppress, localize, or generalize the focal activity, exploring various spatiotemporal dynamics. Traveling waves that emerge from the focus in various phases are investigated and compared with the experimental observations in humans and rats, yielding predictions to be further tested experimentally. The underlying dynamical mechanisms for various phases are also uncovered, using eigenvalue spectra and corresponding eigenmodes at critical states. Finally, we show robustness of the dynamical mechanisms.

### The spatially homogeneous case

Many previous studies of the corticothalamic system investigated its global (spatially uniform) dynamics; e.g., steady states, bifurcations from normal arousal states to epileptic seizures [27, 32, 33]. Specifically, bifurcations have been intensively investigated via changes of the parameter *ν_se_*, and demonstrated that this system can explain primary generalized absence seizures [27, 32]. Furthermore, recent analysis has established a bridge explicitly linking the tractable normal-form dynamical parameters with the underlying physiological ones [33].

In the homogeneous case of generalized absence seizures, the system has spatially uniform activity and the corticothalamic loop experiences a supercritical Hopf bifurcation, then a period-doubling, when *ν_se_* ranging from *ν_se_*(*L*/2) to *ν_se_*(0), as shown in the bifurcation diagram of Fig. 3(A). In Fig. 3(B), various dynamical states are presented by time series of *ϕ_β_* at various *ν_se_*, whose values are correspondingly indicated by the red dashed line in Fig. 3(A). Specifically, one can find oscillating dynamics with a ~ 3 Hz rhythm and the stereotyped SWDs. The presented dynamics can be found over a wide range of parameters. This is one example. This result has been compared with clinical data in detail and thus was employed to explain the pathological transition from normal arousal states to primary generalized absence seizures [27, 32].

**Fig 3.**
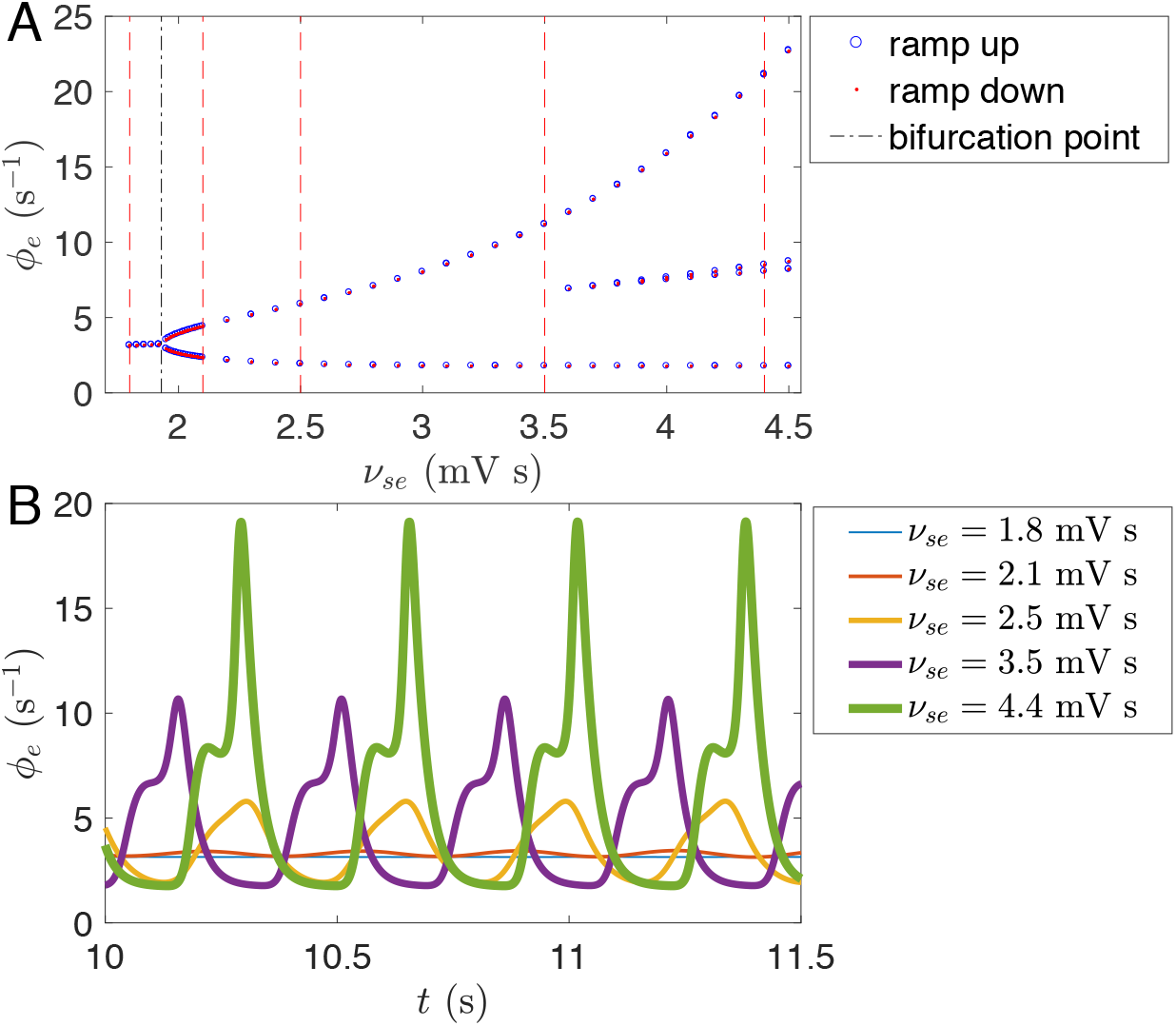
(Color online) Bifurcation in the spatially homogeneous case. (A) Bifurcation diagram of *ϕ_e_* when *ν_se_* ramps up and down from *ν_se_*(*L*/2) to *ν_se_*(0) and vice versa. The black dashed line indicates the threshold 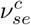 for Hopf bifurcation. (B) Time series of *ϕ_e_* for various *ν_se_* indicated by the red dashed lines in (A).

### From seizure suppression to seizure generalization

Here we show that secondary generalized absence seizures can be induced by the focal activity. The induced absence seizures can have various spatial extents and oscillating frequencies, by exploring the roles of *σ* and *r_e_*. Actually, both of them are constrained by the system size *L*, yielding edge effects. Thus, we investigated various spatiotemporal dynamics vs. *r*_6_/*L* and *σ/L* with *r*_6_/*L* ≪ 1 and *σ/L* ≪ 1, summarized in the phase diagram as shown in Fig. 4, which has six different phases, with Phases I, II, and III discussed in this subsection, and Phases IV, V, and VI in the next.

**Fig 4.**
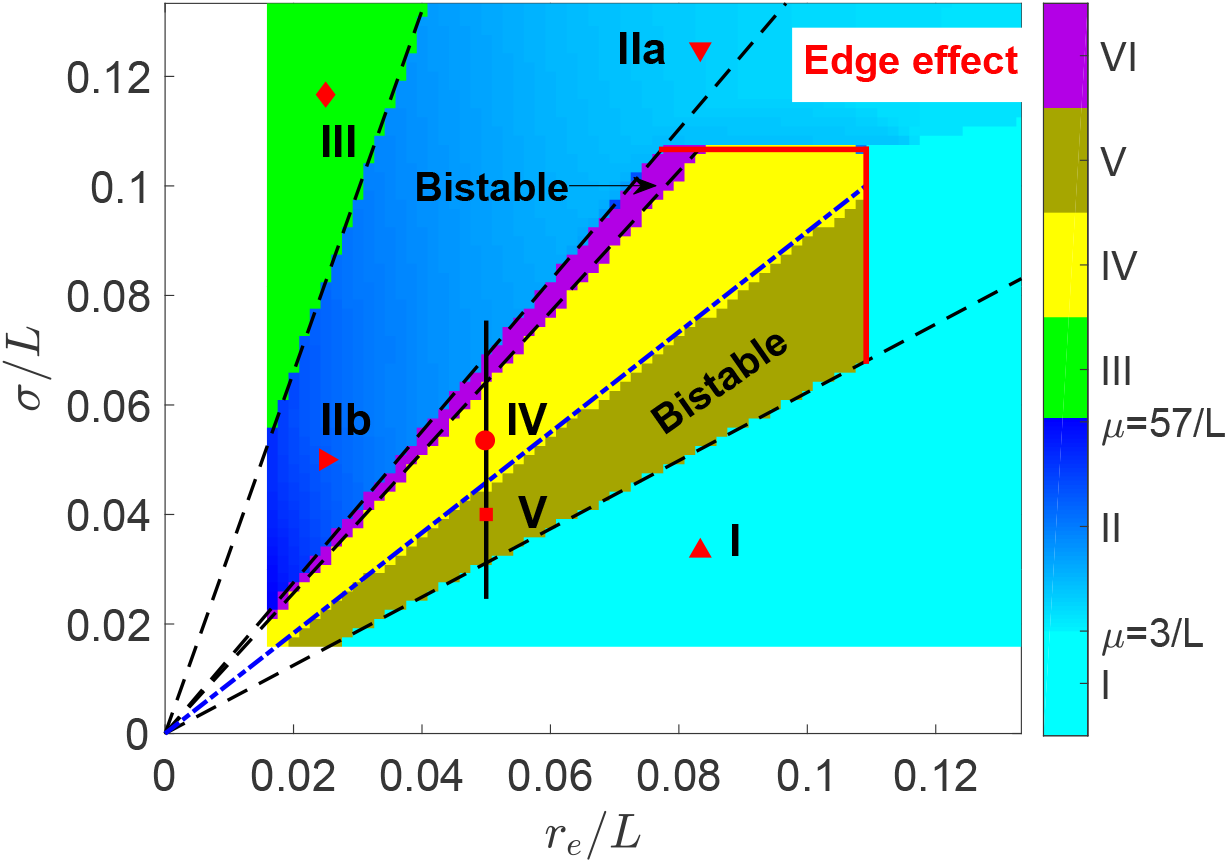
(Color online) Phase diagram of spatiotemporal dynamics vs. axon range *r*_6_ and focal width *σ* both scaled by *L* showing six phases. The instability line for ~ 10 Hz oscillation is predicted by linear stability analysis as indicated the blue dash-dotted line, while other phase separations are indicated by the black dashed lines. Six examples are indicated by various shaped red points, with the corresponding spatiotemporal dynamics of *ϕ*_6_ shown in Fig. 5. The attenuation factor *μ* is also indicated in the colorbar for Phase II, which is scaled by 1/*L*. Note that edge effects limit Phases IV, V, and VI. The black vertical line indicates the parameter range further investigated in Fig. 6. Numerical simulation in the left and bottom white bands would require impractically long simulation time.

In Phase I, the focal activity with small *σ* is suppressed by axonal projections from the normal region if *r_β_* is large as in the case of healthy adults [23]. The activity profile of seizure suppression stabilizes with slightly enhanced activity in the center, as shown in Fig. 5(A).

**Fig 5.**
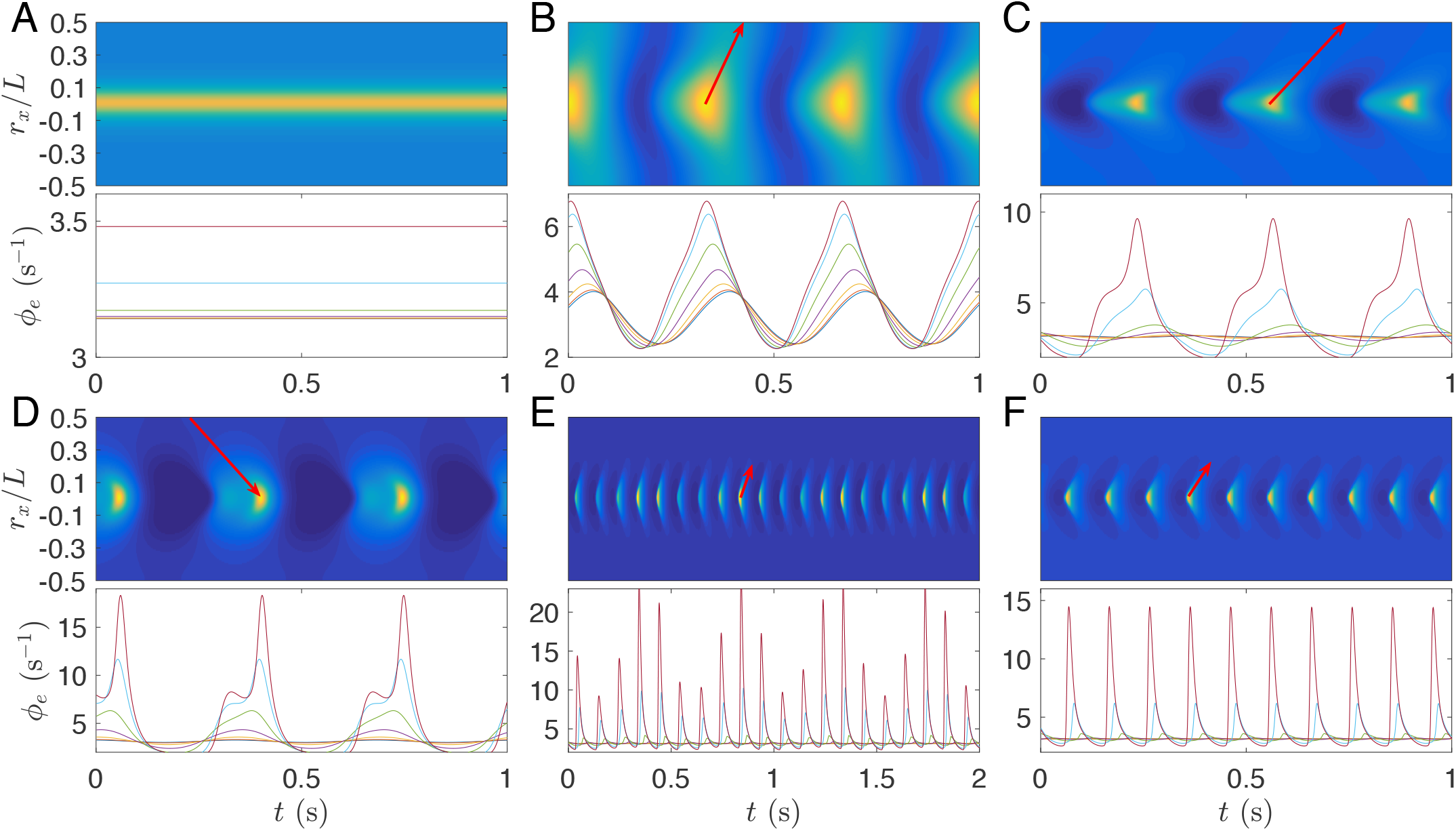
(Color online) Examples of various waves with parameter values of (*r_e_/L,σ/L*) indicated by the corresponding shaped red points in Fig. 4. all dynamics are radially symmetrical, so only 1D wave dynamics is presented. In each case, the upper panel presents the 1D wave dynamics of points at *r_y_* = 0, indicated by the white dashed line in Fig. 2(A), while the lower panel shows the corresponding time series at 7 equispaced points from center to edge with decreasing *ϕ_e_* [for (A)] or its amplitude [for (B) –(F)]. The values *ϕ_e_* in upper panels from dark blue to yellow range from the minimal to the maximal *ϕ_e_* in the corresponding lower panels. (A) Phase I, seizure suppression. (B) Phase IIa, the wave weakly attenuated along *r*. (C) Phase IIb, the wave strongly attenuated along r. (D) Phase III, the wave propagating inward from edge to the center. (E) Phase IV, the spatially localized alpha wave modulated by *a* ~ 2 Hz slow wave. (F) One state in Phase V: the spatially localized regular alpha wave, which coexists with seizure suppression. Wave propagation is indicated by red arrows in (B)-(F), whose slopes are the phase velocities.

In Phase II, the focal activity with large enough *σ* can resist suppression, and even propagate and destabilize the whole system, leading to secondary generalized absence seizures (seizure generalization), which originate from a ~ 3 Hz oscillating focus and propagate rapidly outward to the whole system, but with the wave amplitude attenuated along r, as shown in Figs 5(B) and (C). The attenuation can be characterized by the attenuation factor *μ* = *∂* log *ϕ_e_(r)/∂r*, as indicated in Fig. 4, with larger μ at smaller *r_e_*. Here *μ* is scaled by 1/*L*, with its value characterizing the times of attenuated *ϕ_e_* amplitude when the wave travels through the system. The phase velocity is discussed in the later subsection: Traveling wave properties. Here the system transitions continuously from seizure suppression in Phase I to seizure generalization in Phase II via a supercritical Hopf bifurcation with the oscillating amplitude of *ϕ_e_*(0) increasing gradually from 0, as indicated in the upper right corner of Fig. 4 with boundary induced edge effects.

In Phase III, the direction of propagating waves is reversed from outward in Phase II to inward if *r_e_* is small enough, as seen in Fig. 5(D), with the propagation direction denoted by a red arrow. In future, wavefront instability analysis could be employed to understand such reversal [38].

### Localization of focal activity

In Phase IV, the initial focal activity can induce a localized ~ 10 Hz alpha wave, which decays rapidly in the normal region, as demonstrated in the upper panel of Fig. 5(E), rather than spreading over the whole system as in Figs 5(B)-(D). The alpha wave is not only spatially localized but also temporally modulated at ~ 2 Hz, as shown in Figure 5(E), and reminiscent of complex-partial seizures with impaired consciousness. The complex-partial seizures have stronger 1 - 2 Hz delta-range modulation in the bilateral frontal and parietal neocortex than simple-partial seizures, where consciousness is not impaired [39, 40]. Thus the temporally modulated, spatially localized alpha activity is a potential mechanism for the generation of complex-partial seizures, although we do not consider this point further here.

The above modulation implies the existence of spatiotemporal nonlinear wave interactions. Thus, in Fig. 6, we investigate the bifurcation diagram for the steady state of *ϕ_e_*(0) against *σ/L* at (*r_x_, r_y_*)=(0, 0). It shows that the system experiences a subcritical Hopf bifurcation from a fixed point to a limit cycle, and then a second Hopf bifurcation to a quasiperiodic cycle, e.g., a 2-torus, then a 3-torus, then higher dimensional dynamics, and finally to a chaotic attractor. Such a route to a chaotic attractor is induced by the nonlinear wave interactions, which can also terminate the chaotic dynamics and produce secondary generalized absence seizures when *σ/L* is large enough as shown in the right end of the bifurcation diagram in Fig. 6. This route to a chaotic attractor is different from that observed previously in the homogeneous case, which is induced by the nonlinear corticothalamic interaction loop [33].

**Fig 6.**
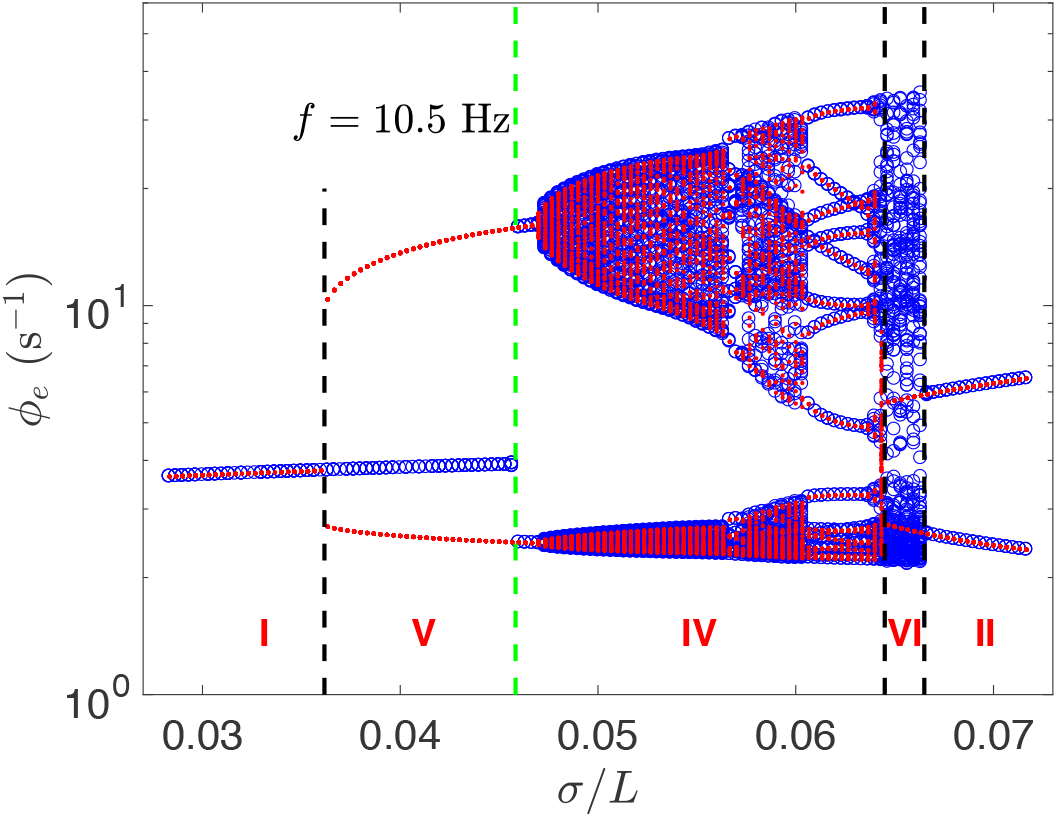
(Color online) Bifurcation diagram of *ϕ_e_*(0) against *σ/L*, with two bistable regions, corresponding to Phase V and VI, respectively. The critical point for bifurcation from seizure suppression to the localized alpha wave with a frequency *f* = 10.5 Hz is consistent with linear stability analysis, as indicated by the green dashed line. Blue circle, ramp up; red dot, ramp down; *r_e_/L* = 0.05.

In Phase V, the system has two stable states: seizure suppression and localized alpha waves (seizure localization). This phase has a smaller *σ/L* than in Phase IV, and a weaker nonlinear wave interaction. As a result, the localized alpha wave here is regular without low frequency modulation, also demonstrated in Fig. 5(F). Actually, such bistability emerges from the subcritical Hopf bifurcation with a hysteresis, and the system experiences a sudden transition from seizure suppression to seizure localization, at the critical point indicated by the green dashed line in Fig. 6. The instability boundary is consistent with the linear stability analysis as introduced in the Methods. This subcritical Hopf bifurcation provides a new route for the transition from normal arousal states to epileptic seizures [6, 41].

In Phase VI, the system has another two stable states: seizure localization and seizure generalization. The value of *σ/L* in this phase is larger than in Phase IV, and there is a stronger nonlinear wave interaction. This bistable region has just a narrow parameter range and may be hard to observe in experiments.

The localized alpha waves in Phases IV, V, and VI are breathers in neural fields [42, 43]. They emerge after seizure suppression becomes unstable and the instability boundary can be well predicted by linear stability analysis, as indicated by the blue dash-dotted line in Fig. 4. All other phase boundaries are also fully determined by the rescaled focal width 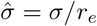 (the dashed lines in Fig. 4). Nonetheless, they terminate at large *σ/L* and *r_e_/L* due to edge effects in the upper right corner of Fig. 4. So seizure localization emerges beyond edge effects, indicating that such phenomenon can exist in general boundary conditions and cortical geometries, e.g., spherical shaped brain.

### Traveling wave properties

From the above, our corticothalamic model with focal spatial heterogeneity can produce secondary generalized absence seizures with ~ 3 Hz traveling waves which spread over the whole system (Phase II), and spatially more localized ~ 10 Hz waves (Phase IV) as observed in the electrophysiological recordings (LFPs, ECoGs, or EEG) in WAG/Rij rats [9]. The spatiotemporal properties in Phase II: seizure generalization and Phase IV: seizure localization, such as spatial extents and temporal frequencies, are comparable with experimental observations in humans and rats, respectively. The resulting cortical waves have robust frequencies in each phase. Therefore, seizure generalization and seizure localization can account for the global and focal aspects of absence epilepsy, respectively, leading to the unification of these two different aspects in our corticothalamic model.

On the other hand, propagating waves of electrical activity in cortex have been observed during seizures in rats, cats, monkeys, and humans, with various wave velocities of 0.01 to 10 m s^−1^, depending on the cortical states and data analysis methods [44–51]. Here we consider the phase velocity v_p_, defined over the whole system for seizure generalization, while for seizure localization, the wave does not propagate over the whole system and we define its effective region to be where oscillating amplitudes are larger than 0.03 times the maximum.

As shown in Fig. 7, for seizure generalization, ~ 3 Hz activity propagates over the whole system at the velocity *υ_p_/L* = 20 to 300 s^−1^, while for seizure localization, ~ 10 Hz activity spreads only within the effective region at *υ_p_/L* = 22 to 60 s^−1^, which is much larger than the axonal propagation speed, which can be estimated to be *υ_e_/L ≈ r_e_γ_e_/L* ≤ 13 s^−1^. In seizure localization, *υ_p_/L* depends linearly on the width *w/L* of the effective region with 0.3 < *w/L* < 0.8, as shown in Fig. 7(A). Figures 7(B) and (C) show that *υ_p_* also depends linearly on *σ* and *r_e_*, with

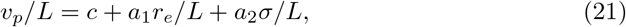

with *c* = 0.80 ± 0.17 s^−1^, *a*_1_ = 440 ± 21 s^−1^ and a_2_ = 107 ± 18 s^−1^. These predictions are potentially testable in experiments.

The linear dependence is consistent with the robust frequency of the localized alpha wave, independent on *r_e_/L* or *σ/L*. As shown in the upper panel of Figs 5(E) and (F) as well as in their eigenmodes to be discussed in the next subsection, the wave has only one peak vs. *r* at each time, yielding the wave number *k* ∝ 1/*w*, and then a constant frequency *f* = *υ_p_k*/2*π*.

**Fig 7.**
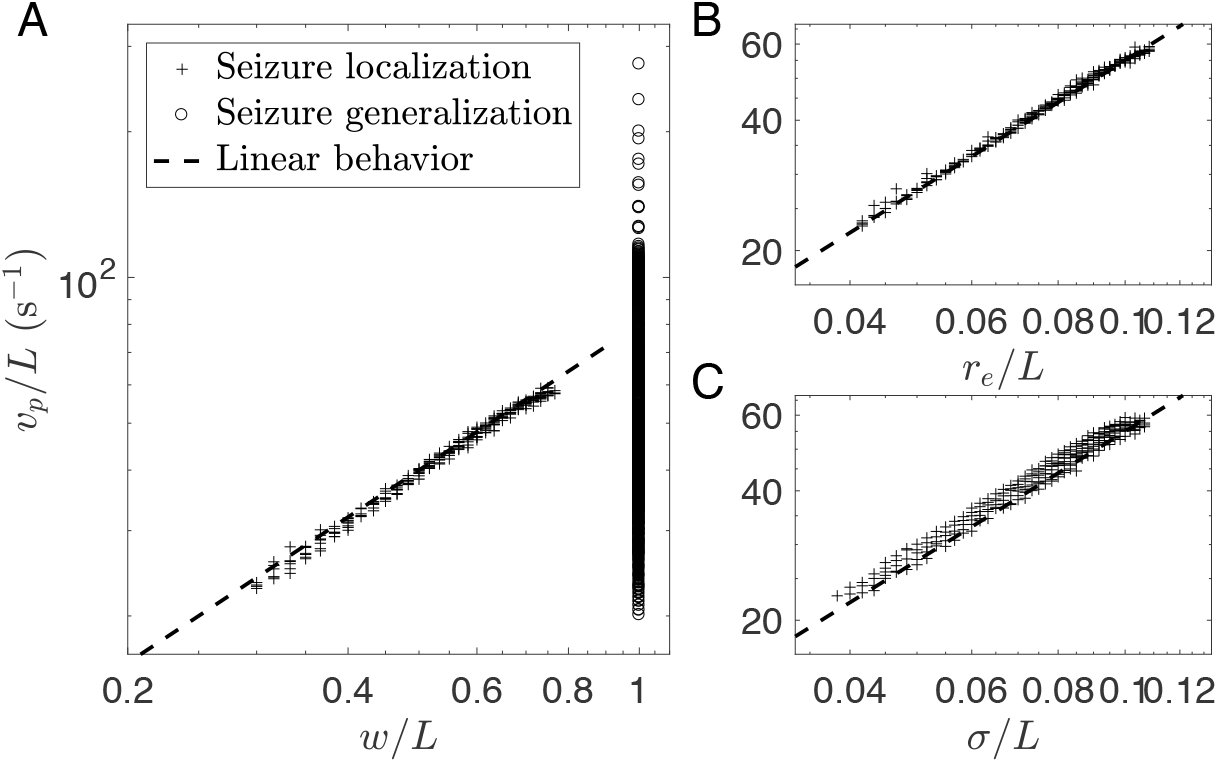
(Color online) Phase velocity *υ_p_* of the traveling waves. The points represent the cases with various parameter pairs (*r_e_/L, σ/L*) in Phases II and IV with *r_e_/L* > 0.05. (A) *υ_p_/L* vs. *w/L* in Phase II (open circle) and Phase IV (plus). (B) *v_p_/L* vs. *r_e_* in Phase IV. (C) *v_p_/L* vs. *σ* in Phase IV. Linear proportionality is indicated by the dashed lines with slope 1 in the log-log plottings.

### Dynamical mechanism underlying emergence

In our previous work, ~ 3 Hz waves and ~ 10 Hz alpha waves were generated due to the resonance of two different delayed feedback loops: *e → r → s → e* or *e → s → e*, respectively, in the homogeneous corticothalamic system [27, 32, 33]. But here a spatially localized alpha wave emerges from the initially focal activity that has an intrinsic ~ 3 Hz rhythm. So, it is not clear whether the localized alpha wave originates from the same resonance of the underlying corticothalamic loop as in the homogeneous system, or is induced by the focal activity via a different dynamical mechanism.

To this end, we investigate the steady states and corresponding eigenvalue spectra of the induced waves, as summarized in Fig. 8. Figures 8(A) and (B) show the three eigenvalues *λ* with largest Re(λ), vs. *σ/L*. The linear stability is determined by the largest Re(λ). In Figs 8(A) and (B), the blue curve corresponds to the localized alpha wave with *f* = 10.5 Hz and the red ones to the generalized waves with *f* = 3.1 Hz.

**Fig 8.**
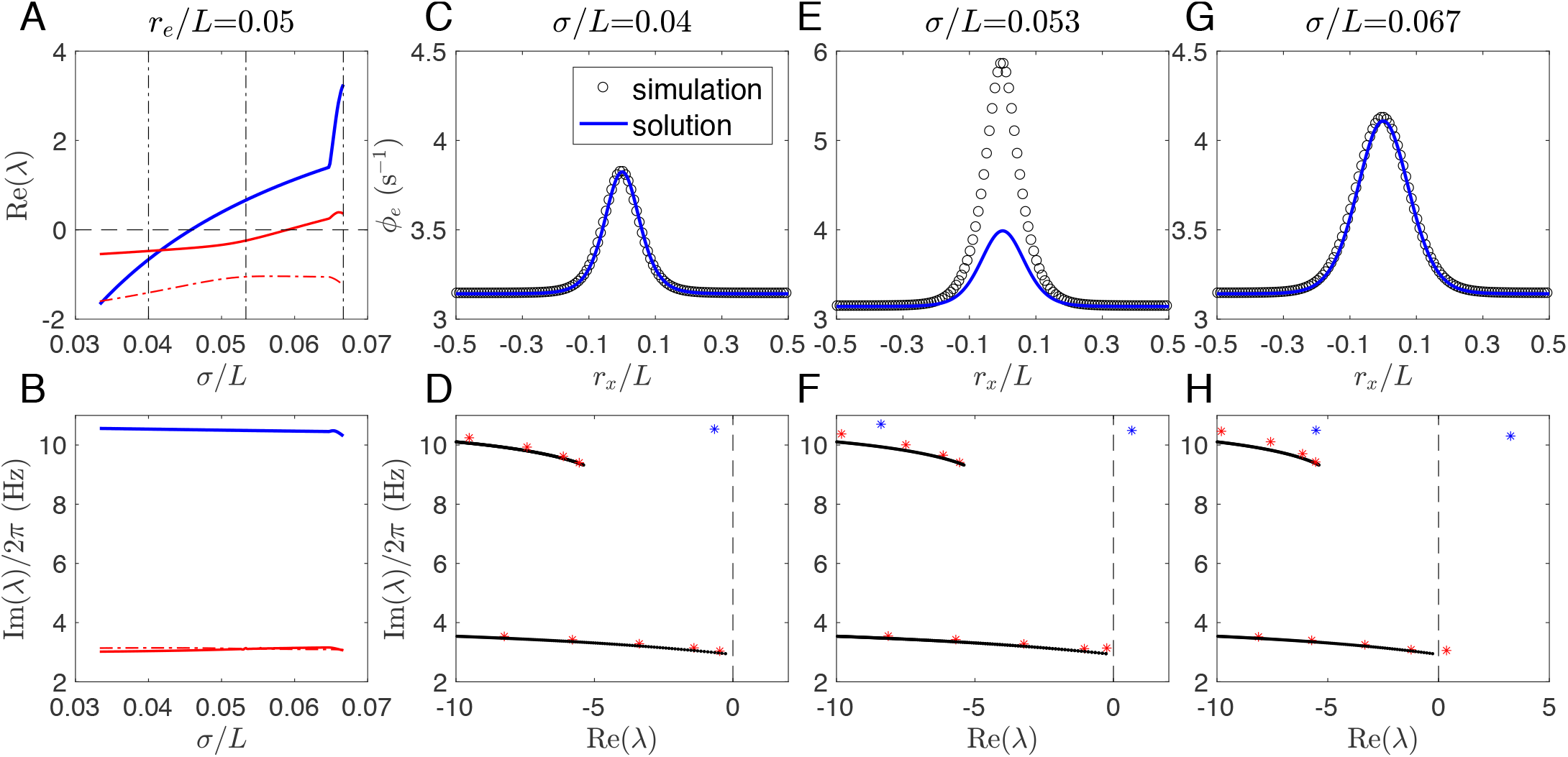
Color online) Steady states and corresponding eigenvalue spectra. (A) Re(λ) vs. *σ/L* at *r_e_/L* = 0.05 for the 3 eigenvalues λ with largest Re(λ). Three vertical dash-dotted black lines indicate the values of *σ/L* for further analysis in (C)-(F). (B) Corresponding Im(λ) vs. *σ/L* as in (A). (C) Stable steady state at *σ/L* = 0.04; (D) Corresponding rightmost eigenvalues (stars) at *σ/L* = 0.04. (E) Unstable steady state evolves to a localized alpha wave at *σ/L* = 0.053; (F) Corresponding rightmost eigenvalues (stars) at *σ/L* = 0.053. (G) Unstable steady state evolves to a ~ 3 Hz wave at *σ/L* = 0.067; (H) Corresponding rightmost eigenvalues (stars) at *σ/L* = 0.067. In (C), (E), and (G), steady states are compared between the time averaged 1D spatial profile from simulations and the numerical solutions obtained from integration of Eq. (10);

Now we focus on the following three typical scenarios: (i) Phase I: Seizure suppression (*σ/L* = 0.04 m); (ii) Phase IV: Seizure localization (*σ/L* = 0.053 m); (iii) Phase II: Seizure generalization (*σ/L* = 0.067 m), with their steady states and corresponding eigenvalue spectra shown in Figs 8(C)–(H). Firstly, for seizure suppression, Fig. 8(C) shows the consistency between the analytical solution and the numerical simulation of the stable steady state, with its stability indicated in Fig. 8(D). Their excellent match verifies the assumption of radial symmetry, and lends confidence in further analysis. Secondly, for seizure localization with larger *σ/L*, the steady state becomes unstable with mismatch between the analytical solution and the numerical simulation in the focal region, as shown in Fig. 8(E). Its instability is indicated by one eigenvalue with a positive real part [the rightmost blue star in Fig. 8(F)]. Here, the focal acitivity is localized without spreading over the whole system because *σ/L* is only large enough to free the focal activity from being suppressed by the normal region. So the steady state is stable at the tail but unstable in the focal region. Finally, for seizure generalization with *σ/L* further increased, the steady state at the tail becomes unstable as well, and a ~ 3 Hz wave originating from the focal region propagates over the whole system. One can find from Fig. 8(H) that there is one more eigenvalue along the bottom branch crossing the imaginary axis. Nonetheless, Fig. 8(G) shows the consistency between the time-averaged activity and the unstable steady state, suggesting that the activity at each spatial point surrounds the unstable steady state, and the ~ 3 Hz wave emerges via a supercritical Hopf bifurcation.

The dynamical picture is consistent with that indicated in eigenvalue spectra. Figures 8(D), (F), and (H) show that most eigenvalues (red stars) align with two branches (black lines), where 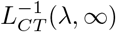 is real negative 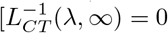 at the branch right end]. These two branches correspond to the resonance of two delayed feedback loops: *e → r → s → e* or *e → s → e*, respectively, as in the spatially homogeneous case [27, 33]. It is a general property in the delayed system that an infinite number of eigenvalues will align with such branches. Generally, the delay leads the spectrum of the system to be determined by the solutions of

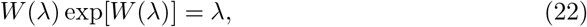

where *W*(λ) is a complex function with an infinite number of solutions [52]. So here most dynamical eigenmodes along the two branches correspond to the potential resonance of the same corticothalamic loops as the spatially homogeneous case.

However, some eigenvalues [blue stars in Figs 8(D), (F), and (H)] emerge beyond the last two branches, implying that their eigenmodes are not purely induced by the resonance of the underlying corticothalamic loops. For these eigenmodes, 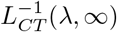 has a much larger value than those with eigenvalues (red stars) close to the two branches. That means the steady state at the tail stays stable and far away from a corticothalamic loop resonance, while the system’s steady state is unstable, as indicated in Fig. 8(F) with a positive Re(λ). Thus, this eigenmode is confined in the focal region, as confirmed in Figs 9(A) and (B). As a result, it emerges a different spatiotemporal wave with different frequency and waveform from that in seizure generalization. What is more, this eigenmode is also spatially confined by its steady state, as shown in Fig. 9(B). This is in contrast with the eigenmode in seizure generalization, as shown in Figs 9(C) and (D), where the wave propagates outwards from the focal area, but the activity at each spatial point still surrounds its steady state.

**Fig 9.**
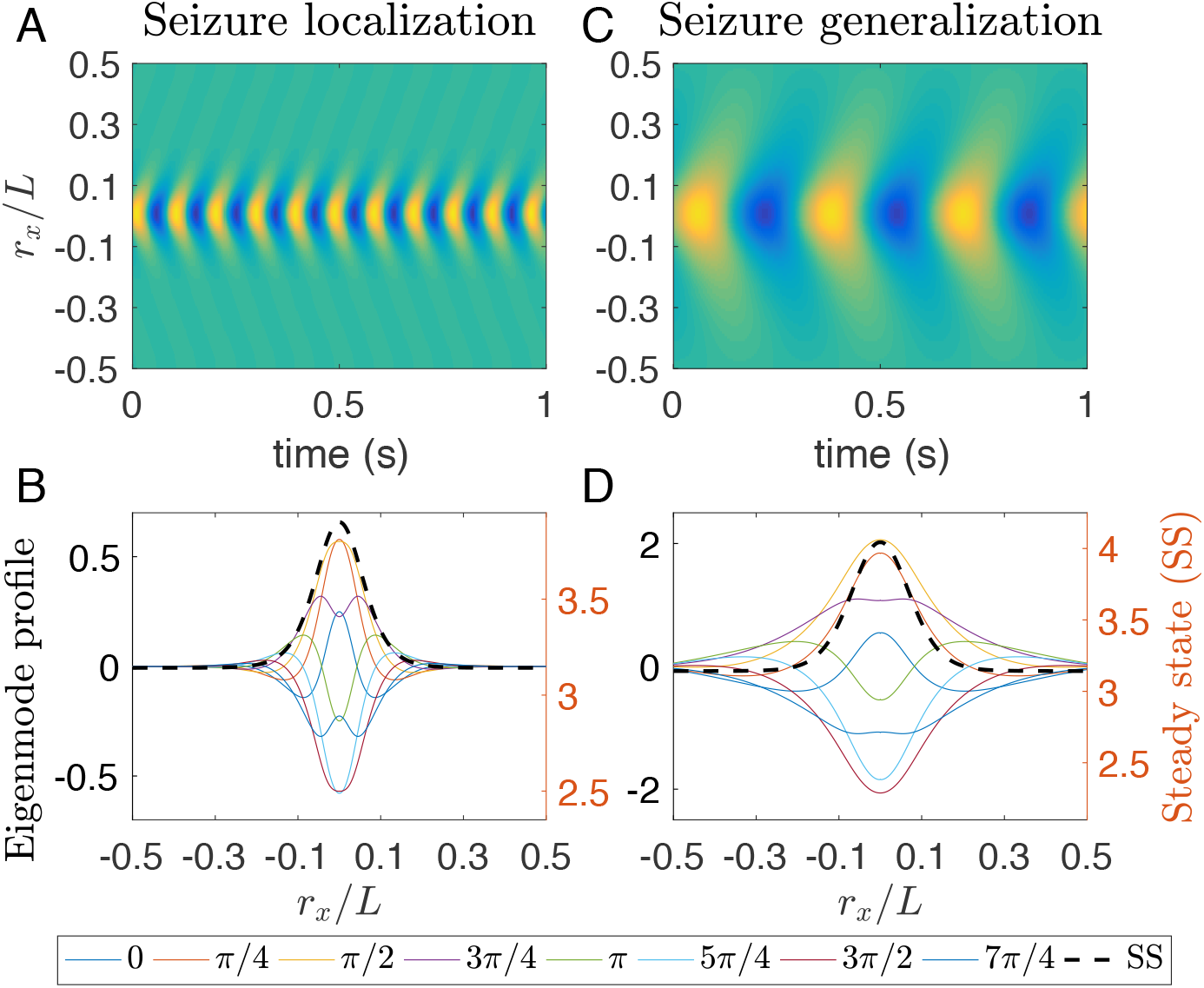
(Color online) Spatiotemporal eigenmodes of ~ 10 Hz wave and ~ 3 Hz wave at the respective critical points for *r_e_/L* = 0.05. (A) 1D wave dynamics of *ϕ_e_* for points at *r_y_* = 0 with *σ/L* = 0.046 for seizure localization. (B) 1D spatial profile of *ϕ_e_* with *σ/L* = 0.046 for both eigenmodes and the corresponding steady states at *r_y_* = 0 and various instantaneous phases *φ*_0_, as indicated in the bottom legend. (C) same as (A) but with *σ/L* = 0.059 for seizure generalization. (D) same as (B) but with *σ/L* = 0.059.

Thus, the localized alpha waves emerge due to spatial confinement, which is different from the corticothalamic loop resonance in the spatially homogeneous case [27, 32, 33]. This mechanism also explains why edge effects prevent seizure localization when there is not enough space to confine the wave away from the boundary. Furthermore, the spatially confined eigenmode in Figs 9(A) and (B) is also unstable and then further evolves to another localized alpha wave with a much higher amplitude, as seen in Fig. 8(E). So seizure localization emerges suddenly from seizure suppression via a subcritical Hopf bifurcation.

### Robustness of dynamical mechanism

The above dynamical mechanism for seizure localization is robust and does not require parameter fine-tuning. A large parameter region can be found in Figs 10(A) and (B), where the focal activity is too unstable to be suppressed by the normal region, but doesn’t generalize over the whole system, with *ν_se_*(0) much larger than the linear stability boundary of the corticothalamic loops [the critical value 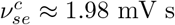, as indicated in Fig. 2(A) and Fig. 3(A)]. Figure 10(A) also shows that seizure suppression can transition directly to seizure generalization, when *ν_se_*(0) is not large enough to induce seizure localization. This is the weakly nonlinear case explored in [53]. So seizure localization emerges due to the combination of nonlinear effects and spatial comfinement.

**Fig 10.**
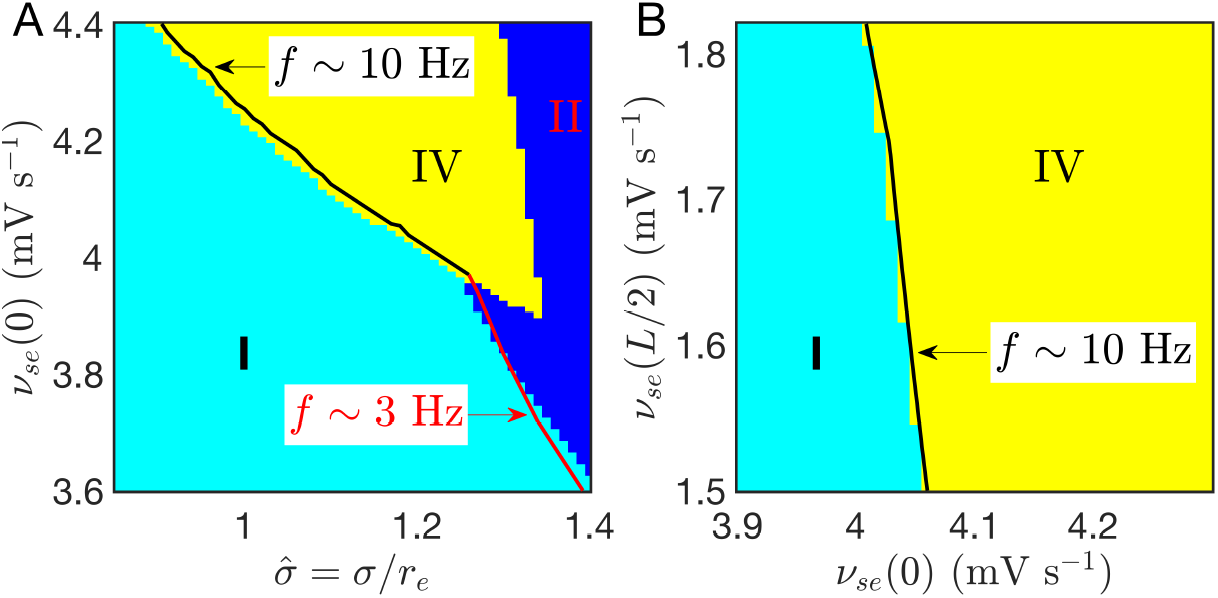
(Color online) Seizure localization due to strong nonlinear effects of the focal activity. (A) Phase diagram on 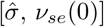; (B) Phase diagram on [*ν_se_* (0), *ν_se_* (*L*/2)] with *σ* = 1.2. *ν_se_*(*L*/2) has a much weaker effect on the emergence of seizure localization, in consistent with its underlying dynamical mechanism. The linear instability boundaries are also well predicted by linear stability analysis.

The emergence also requires the normal region to be stable enough to be unevoked by the focal activity. However, Fig. 10(B) shows that *ν_se_*(*L*/2) has a much weaker effect. This is consistent with the underlying dynamical mechanism: for seizure localization, the normal region is already stable and can resist invasion of seizure activity of the seizure focus, so further lowering *ν_se_*(*L*/2) to render the normal region more stable will make little contribution, as long as *ν_se_*(0) is large enough to free the focal region from being suppressed by the normal region. These results are also consistent with the experimental findings in WAG/Rij and GAERS rats: Pharmacological deactivation by blocking the neural activity of the driving cortical source can almost completely abolish SWDs in all cortical regions [17], while such deactivation in other cortical regions has little effect [54].

The simulation results are also confirmed by linear stability analysis, as shown in Fig. 10, indicating that the parameter selection for the dynamical mechanism is not unique, but can be general, whenever the system has strong nonlinear effects in the focal area and the conditions for spatial comfinement are satisfied.

## Summary and Discussion

This work has investigated suppression, localization, and generalization of focal activity, via a physiology-based corticothalamic model with focal spatial heterogeneity. We found that the interplay between cortical propagation and the underlying corticothalamic circuit can generate various spatiotemporal dynamics, which mainly depend on the focal width *σ* and the axon range *r_e_* scaled by the system size *L*. The main results are:

i. The spatiotemporal dynamic summarized in the phase diagram (Fig. 4) has 6 phases, which can be categorized into three scenarios: suppression (Phase I and V), generalization (Phases II, III, and VI), and localization (Phases IV, V, and VI) of focal activity.
ii. Axonal projections from the normal region can suppress the focal activity when 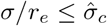, whose value is mainly determined by the pathological value *ν_se_*(0) of focal corticothalamic connection strength in this study.
iii. In seizure generalization, initial ~ 3 Hz focal activity propagates rapidly over the whole system, with the temporal frequency and the spatial extent comparable with absence seizure activity in humans, and can be associated with the global aspect of absence epilepsy. Besides, the direction of propagating wave can be reversed from outward to inward if *r_e_/L* is small.
iv. In seizure localization, spatially localized ~ 10 Hz activity emerges due to strong nonlinear effects in the focal region and spatial comfinement by the surrounding stable normal region, with the temporal frequency and the spatial extent comparable with absence seizure activity in genetic rat models, and can be associated with the focal aspect of absence epilepsy. This also provides a biophysical explanation of spatially more localized waves with higher oscillating frequency observed in such rats. Besides, its emergence is beyond edge effects, and thus general for other boundary conditions and cortical geometries.
v. In both seizure generalization and seizure localization, the oscillating frequencies are robust, independent on *r_e_/L* and *σ/L*, and the phase velocity *υ_p_* of the waves are much larger than the axonal propagation speed.
vi. In seizure localization, *υ_p_* depends linearly on the spatial extent *w*, which can be further tested in experiments.
vii. The underlying dynamical mechanisms for both seizure generalization and seizure localizatoin are explained in detail through eigenvalue spectra and corresponding eigenmodes at critical states.
viii. The state in the normal region has a weaker effect on suppressing the focal activity than the state in the focal region. This result is consistent with the experimental findings of the effects of pharmacological deactivations on different cortical regions in WAG/Rij and GAERS rats [17, 54].

Our results also present several predictions for further experimental test:

i. The 7 – 11 Hz activity in WAG/Rij and GAERS rats may be explained by a relative scaled short axon length *r_e_/L*, while *r_e_/L* in human brains may be large enough to place the system in the regime dominated by edge effects, and therefore induce seizure generalization with the focal activity spreading over the whole system.
ii. The localized alpha waves in seizure localization can be modulated by ~ 2 Hz slow waves. It is worth further investigating the underlying biophysical mechanism for such modulation with robust ~ 2 Hz frequency in future.
iii. In a narrow parameter range, the system can have two stable states: seizure localization and seizure generalization, referred as the Phase VI shown in Fig. 4 and Fig. 6. This phenomenon may explain the on-off intermittency of SWDs durations of absence seizures in the EEG of WAG/Rij rats [55], which also has chaotic properties.

In this work, we have focused on the interplay of cortical propagation and the underlying corticothalamic loop, which goes beyond most previous work about spatially extended heterogeneous but purely cortical models [56–59]. The emergence of breathers during seizure localization was explored in previous work, but only in an abstract neural field model with Mexican hat connection profile and feedbacks, such as spike-frequency adaptation [42, 60, 61]. In most previous theoretical work, the sigmoid function was simplified to a step function, easing linear stability analysis [38, 62]; however, the step function is unrealistic, whereas our corticothalamic model is physiologically justifiable, and relies on parameters that are experimentally measurable.

Further extensions should be investigated in future to account for more features of absence seizure activities; Some include:

i. Thalamic relay neurons have well-characterized dual firing modes: bursting and tonic spiking. It was found recently in experiments that the rhythmic synchronized phasic firing of thalamic relay neuronal population can initiate SWDs and seems necessary for absence seizure maintenance [19]. The origin of such synchronization is shown from cortical drive and temporally framing via feedforward inhibition from cortical neurons to reticular neurons and then to relay neurons [20]. So it is essential to study the effect of neuronal firing modes and its interaction with the corticothalamic loop, to bridge the gap between microscale mechanisms and macroscopic cortico-thalamo-cortical oscillations in absence epilepsy.
ii. It has been widely explored how SWDs can be more robustly produced by involving a second slow inhibitory population or inhibitory variable with a slower time scale, such as GABA_*B*_ [5, 63]. The relative contribution of both fast GABA_*A*_ and slow GABA_*B*_ receptor-mediated inhibition has been discussed for controlling absence seizures [5, 64]. The slow GABA_*B*_ dynamics should be included to account for the detailed wave shape of SWDs.
iii. The specific somatosensory-thalamo-cortical network should be considered to further study the dynamical mechanism for SWD initiation, maintenance, and termination in genetic rat models [18, 65]. The exact interactions between cortex and different thalamic nuclei are essential for understanding how the cortical ‘focus’ within the perioral subregion of the somatosensory cortex drives other parts of the cortex and the ventral basal complex of the thalamus.
iv. Lesions and pharmacological manipulations of the basal ganglia (BG) and neuromodulator pathways affect SWDs [66, 67]. The effect of BG has been proposed to be important for maintaining absence seizures over several tens of seconds by the dynamical loop of the BG-thalamo-cortical network [68], which also provides a modulation site and an intervention pathway to prevent absence seizures [69, 70]. So it is essential to investigate the effect of BG modulation on the suppression, localization, and generalization of focal activity.
v. The transitions from seizure suppression to seizure localization or seizure generalization can be more systematically studied by employing normal form theory [33]. However, the canonical nature in the sense of normal form theory has not yet been rigorously justified for spatiotemporal evolution equations. But with some special perturbation modes, it can still be employed to understand the effects of other biophysical properties on the bifurcation types, such as bursting activities in thalamic populations, the slow dynamics of extracellular potassium ion concentration, and spike frequency adaptation.

## Author Contributions

Conceived and designed research: D.-P.Y, P.A.R. Performed research: D.-P.Y, P.A.R. Wrote the paper: D.-P.Y, P.A.R.

## Acknowledgments

This work was supported by the Australian Research Council Center for Integrative Brain Function (Grant CE140100007) and by Australian Research Council Laureate Fellowship Grant FL140100025.

## A Derivation of *L_CT_* (λ,*r*)

Here we present the derivation of the radially symmetric linear operator *L_CT_*(λ, *r*) for each spatial point of the corticothalamic system and each eigenmode corresponding to eigenvalue λ. The eigenvalue λ = 0 is for the stationary solution in Eqs (10) and (12) and others for general eigenmodes in Eqs (18) and (20).

We derive *L_CT_*(λ, *r*) for both cases by imposing a small perturbation on either 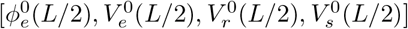 or 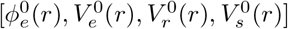. Here the general perturbation with eigenvalue λ at position *r* is denoted as 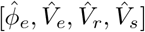, where (λ, *r*) is ignored for simplicity, which yields

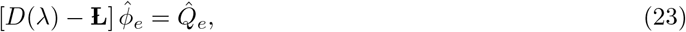

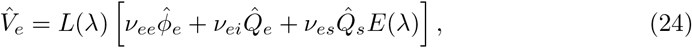

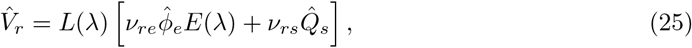

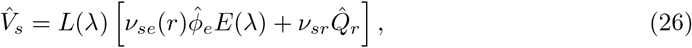

with

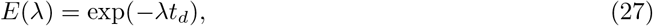

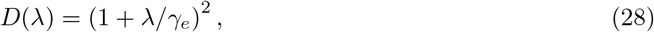

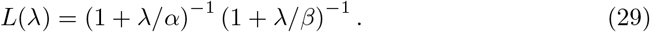

By linearizing the sigmoid function near the steady state, Eqs (24)–(26) yield:

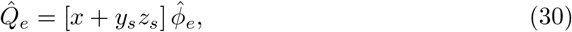

with

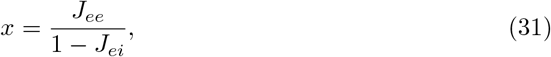

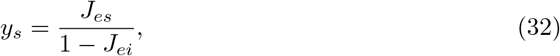

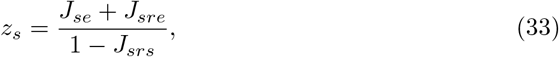

where *J_abc_* = *J_ab_J_bc_*, and *J_ab_*(λ, *r*) = *G_ab_(r)L*(λ) for connections within cortex or thalamus, and *J_ab_*(λ, *r*) = *G_ab_(r)L*(λ)*E*(λ) for connections between cortex and thalamus with time delay *t_d_*. Here *x*, *y_s_*, and *z_s_* are the signal propagations along axons from cortical field to cortex, from thalamic relay nucleus to cortex, and from cortical field to thalamic relay nucleus, respectively, as shown in Fig. 2 of Reference [33]. The gain *G_ab_(r)* = *ν_ab_η_a_(r)Q*_max_/*σ′* is the additional output produced by neurons a per unit additional input from neurons b. Using Eqs (23)–(33), *L_CT_*(λ, *r*) can be expressed as

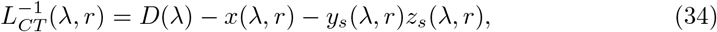

which at 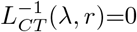 yields the resonance of the corticothalamic loop for each radial distance *r* and each eigenmode with eigenvalue λ.

## References

1. Crunelli V, Leresche N. Childhood absence epilepsy: Genes, channels, neurons and networks. Nat Rev Neuroscix. 2002;3:371.

2. Meeren HKM, van Luijtelaar G, Lopes da Silva FH, Coenen AML. Evolving concepts on the pathophysiology of absence seizures: The cortical focus theory. JAMA Neurol. 2005;62(3):371–376.

3. Depaulis A, Charpier S. Pathophysiology of absence epilepsy: Insights from genetic models. Neurosci Lett. 2018;667:53–65.

4. Fisher RS, Prince DA. Spike-wave rhythms in cat cortex induced by parenteral penicillin. I. Electroencephalographic features. Electroencephalogr Clin Neurophysiol. 1977;42(5):608–624.

5. Destexhe A. Spike-and-Wave Oscillations Based on the Properties of GABAB Receptors. J Neurosci. 1998;18(21):9099–9111.

6. Lopes da Silva FH, Blanes W, Kalitzin SN, Parra J, Suffczynski P, Velis DN. Epilepsies as Dynamical Diseases of Brain Systems: Basic Models of the Transition Between Normal and Epileptic Activity. Epilepsia. 2003;44(s12):72–83.

7. Meeren HKM, Veening JG, Moderscheim TAE, Coenen AML, van Luijtelaar G. Thalamic lesions in a genetic rat model of absence epilepsy: Dissociation between spike-wave discharges and sleep spindles. Exp Neurol. 2009;217(1):25–37.

8. Avoli M. A brief history on the oscillating roles of thalamus and cortex in absence seizures. Epilepsia. 2012;53(5):779–789.

9. Meeren HKM, Pijn JPM, van Luijtelaar ELJM, Coenen AML, Lopes da Silva FH. Cortical Focus Drives Widespread Corticothalamic Networks during Spontaneous Absence Seizures in Rats. J Neurosci. 2002;22(4):1480–1495.

10. Sitnikova E, van Luijtelaar G. Cortical control of generalized absence seizures: effect of lidocaine applied to the somatosensory cortex in WAG/Rij rats. Brain Res. 2004;1012(1):127–137.

11. Polack PO, Guillemain I, Hu E, Deransart C, Depaulis A, Charpier S. Deep Layer Somatosensory Cortical Neurons Initiate Spike-and-Wave Discharges in a Genetic Model of Absence Seizures. J Neurosci. 2007;27(24):6590–6599.

12. Polack PO, Mahon S, Chavez M, Charpier S. Inactivation of the Somatosensory Cortex Prevents Paroxysmal Oscillations in Cortical and Related Thalamic Neurons in a Genetic Model of Absence Epilepsy. Cereb Cortex. 2009;19(9):2078–2091.

13. Scicchitano F, van Rijn CM, van Luijtelaar G. Unilateral and Bilateral Cortical Resection: Effects on Spike-Wave Discharges in a Genetic Absence Epilepsy Model. PLOS ONE. 2015;10(8):1–20.

14. Holmes MD, Brown M, Tucker DM. Are “Generalized” Seizures Truly Generalized? Evidence of Localized Mesial Frontal and Frontopolar Discharges in Absence. Epilepsia. 2004;45(12):1568–1579.

15. Sarrigiannis PG, Zhao Y, He F, Billings SA, Baster K, Rittey C, et al. The cortical focus in childhood absence epilepsy; evidence from nonlinear analysis of scalp EEG recordings. Clin Neurophysiol. 2018;129(3):602–617.

16. Blumenfeld H. Cellular and Network Mechanisms of Spike-Wave Seizures. Epilepsia. 2005;46(s9):21–33.

17. van Luijtelaar G, Sitnikova E. Global and focal aspects of absence epilepsy: The contribution of genetic models. Neurosci Biobehav Rev. 2006;30(7):983–1003.

18. Lüttjohann A, van Luijtelaar G. Dynamics of networks during absence seizure’s on- and offset in rodents and man. Front Physiol. 2015;6:16.

19. Sorokin JM, Davidson TJ, Frechette E, Abramian AM, Deisseroth K, Huguenard JR, et al. Bidirectional Control of Generalized Epilepsy Networks via Rapid Real-Time Switching of Firing Mode. Neuron. 2017;93(1):194–210.

20. McCafferty C, David F, Venzi M, Lorincz ML, Delicata F, Atherton Z, et al. Cortical drive and thalamic feed-forward inhibition control thalamic output synchrony during absence seizures. Nat Neurosci. 2018;21(5):744–756.

21. Coenen AML, van Luijtelaar ELJM. Genetic Animal Models for Absence Epilepsy: A Review of the WAG/Rij Strain of Rats. Behav Genet. 2003;33(6):635–655.

22. Neubauer FB, Sederberg A, MacLean JN. Local changes in neocortical circuit dynamics coincide with the spread of seizures to thalamus in a model of epilepsy. Front Neural Circuits. 2014;8:101.

23. Robinson PA, Rennie CJ, Rowe DL, O’Connor SC. Estimation of multiscale neurophysiologic parameters by electroencephalographic means. Hum Brain Mapp. 2004;23(1):53–72.

24. Nersesyan H, Hyder F, Rothman DL, Blumenfeld H. Dynamic fMRI and EEG Recordings during Spike-Wave Seizures and Generalized Tonic-Clonic Seizures in WAG/Rij Rats. J Cereb Blood Flow Metab. 2004;24(6):589–599.

25. Lopes da Silva FH, van Rotterdam A, van Leeuwen WS, Tielen AM. Dynamic characteristics of visual evoked potentials in the dog. II. Beta frequency selectivity in evoked potentials and background activity. Electroencephalogr Clin Neurophysiol. 1970;29(3):260–268.

26. Robinson PA. Neurophysical theory of coherence and correlations of electroencephalographic and electrocorticographic signals. J Theor Biol. 2003;222(2):163–175.

27. Robinson PA, Rennie CJ, Rowe DL. Dynamics of large-scale brain activity in normal arousal states and epileptic seizures. Phys Rev E. 2002;65:041924.

28. Jirsa VK, Haken H. Field Theory of Electromagnetic Brain Activity. Phys Rev Lett. 1996;77:960–963.

29. Robinson PA, Rennie CJ, Wright JJ. Propagation and stability of waves of electrical activity in the cerebral cortex. Phys Rev E. 1997;56(1):826.

30. Braitenberg V, Schüz A. Cortex: statistics and geometry of neuronal connectivity. Springer Science & Business Media; 2013.

31. Wright JJ, Liley DTJ. Dynamics of the brain at global and microscopic scales: Neural networks and the EEG. Behav Brain Sci. 1996;19(02):285–295.

32. Breakspear M, Roberts JA, Terry JR, Rodrigues S, Mahant N, Robinson PA. A unifying explanation of primary generalized seizures through nonlinear brain modeling and bifurcation analysis. Cereb Cortex. 2006;16(9):1296–1313.

33. Yang DP, Robinson PA. Critical dynamics of Hopf bifurcations in the corticothalamic system: Transitions from normal arousal states to epileptic seizures. Phys Rev E. 2017;95:042410.

34. Destexhe A, Sejnowski TJ. Thalamocortical Assemblies. Oxford: Oxford University Press; 2001.

35. McCormick DA, Contreras D. On The Cellular and Network Bases of Epileptic Seizures. Annu Rev Physiol. 2001;63(1):815–846.

36. Sanz-Leon P, Robinson PA, Knock SA, Drysdale PD, Abeysuriya RG, Fung PK, et al. NFTsim: Theory and Simulation of Multiscale Neural Field Dynamics. bioRxiv. 2018;.

37. Firth WJ, Scroggie AJ. Optical Bullet Holes: Robust Controllable Localized States of a Nonlinear Cavity. Phys Rev Lett. 1996;76:1623–1626.

38. Coombes S, Owen MR. Evans Functions for Integral Neural Field Equations with Heaviside Firing Rate Function. SIAM J Appl Dyn Syst. 2004;3(4):574–600.

39. Englot DJ, et al. Impaired consciousness in temporal lobe seizures: role of cortical slow activity. Brain. 2010;133(12):3764–3777.

40. Rosanova M, Gosseries O, Casarotto S, Boly M, Casali AG, Bruno MA, et al. Recovery of cortical effective connectivity and recovery of consciousness in vegetative patients. Brain. 2012;135(4):1308–1320.

41. Lopes da Silva FH, Blanes W, Kalitzin SN, Parra J, Suffczynski P, Velis DN. Dynamical diseases of brain systems: different routes to epileptic seizures. IEEE Trans Biomed Eng. 2003;50(5):540–548.

42. Folias SE, Bressloff PC. Breathers in Two-Dimensional Neural Media. Phys Rev Lett. 2005;95:208107.

43. Folias SE. Traveling waves and breathers in an excitatory-inhibitory neural field. Phys Rev E. 2017;95:032210.

44. Chervin RD, Pierce PA, Connors BW. Periodicity and directionality in the propagation of epileptiform discharges across neocortex. J Neurophysiol. 1988;60(5):1695–1713.

45. Kramer MA, Kirsch HE, Szeri AJ. Pathological pattern formation and cortical propagation of epileptic seizures. J Royal Soc Interface. 2005;2(2):113–127.

46. Pinto DJ, Patrick SL, Huang WC, Connors BW. Initiation, Propagation, and Termination of Epileptiform Activity in Rodent Neocortex In Vitro Involve Distinct Mechanisms. J Neurosci. 2005;25(36):8131–8140.

47. Trevelyan AJ, Sussillo D, Yuste R. Feedforward Inhibition Contributes to the Control of Epileptiform Propagation Speed. J Neurosci. 2007;27(13):3383–3387.

48. Muller L, Destexhe A. Propagating waves in thalamus, cortex and the thalamocortical system: Experiments and models. Journal of Physiology-Paris. 2012;106(5):222–238. New trends in neurogeometrical approaches to the brain and mind problem.

49. Trevelyan AJ, Schevon CA. How inhibition influences seizure propagation. Neuropharmacology. 2013;69:45–54. New Targets and Approaches to the Treatment of Epilepsy.

50. Martinet LE, Ahmed OJ, Lepage KQ, Cash SS, Kramer MA. Slow Spatial Recruitment of Neocortex during Secondarily Generalized Seizures and Its Relation to Surgical Outcome. J Neurosci. 2015;35(25):9477–9490.

51. Martinet LE, Fiddyment G, Madsen JR, Eskandar EN, Truccolo W, Eden UT, et al. Human seizures couple across spatial scales through travelling wave dynamics. Nat Commun. 2017;8:14896.

52. Grindrod P, Pinotsis DA. On the spectra of certain integro-differential-delay problems with applications in neurodynamics. Physica D. 2011;240(1):13–20.

53. Kim JW, Roberts JA, Robinson PA. Dynamics of epileptic seizures: Evolution, spreading, and suppression. J Theor Biol. 2009;257(4):527–532.

54. van Luijtelaar G, Sitnikova E, Luttjohann A. On the Origin and Suddenness of Absences in Genetic Absence Models. Clin EEG Neurosci. 2011;42(2):83–97.

55. Sitnikova E, Hramov AE, Grubov VV, Ovchinnkov AA, Koronovsky AA. On-off intermittency of thalamo-cortical oscillations in the electroencephalogram of rats with genetic predisposition to absence epilepsy. Brain Res. 2012;1436:147–156.

56. Taylor PN, Baier G. A spatially extended model for macroscopic spike-wave discharges. J Comput Neurosci. 2011;31(3):679–684.

57. Goodfellow M, Schindler K, Baier G. Intermittent spike-wave dynamics in a heterogeneous, spatially extended neural mass model. NeuroImage. 2011;55(3):920–932.

58. Marc G, Neal TP, Yujiang W, James GD, Gerold B. Modelling the role of tissue heterogeneity in epileptic rhythms. Eur J Neurosci. 2012;36(2):2178–2187.

59. Taylor PN, Goodfellow M, Wang Y, Baier G. Towards a large-scale model of patient-specific epileptic spike-wave discharges. Biol Cybern. 2013;107(1):83–94.

60. Coombes S, Owen MR. Bumps, Breathers, and Waves in a Neural Network with Spike Frequency Adaptation. Phys Rev Lett. 2005;94:148102.

61. Qi Y, Gong P. Dynamic patterns in a two-dimensional neural field with refractoriness. Phys Rev E. 2015;92:022702.

62. Loxley PN, Robinson PA. Soliton Model of Competitive Neural Dynamics during Binocular Rivalry. Phys Rev Lett. 2009;102:258701.

63. Marten F, Rodrigues S, Suffczynski P, Richardson MP, Terry JR. Derivation and analysis of an ordinary differential equation mean-field model for studying clinically recorded epilepsy dynamics. Phys Rev E. 2009;79:021911.

64. Chen M, Guo D, Xia Y, Yao D. Control of Absence Seizures by the Thalamic Feed-Forward Inhibition. Front Comput Neurosci. 2017;11:31.

65. Sysoeva MV, Lüttjohann A, van Luijtelaar G, Sysoev IV. Dynamics of directional coupling underlying spike-wave discharges. Neuroscience. 2016;314:75–89.

66. Danober L, Depaulis A, Marescaux C, Vergnes M. Effects of cholinergic drugs on genetic absence seizures in rats. Eur J Pharmacol. 1993;234(2):263–268.

67. Danober L, Deransart C, Depaulis A, Vergnes M, Marescaux C. Pathophysiological mechanisms of genetic absence epilepsy in the rat. Prog Neurobiol. 1998;55(1):27–57.

68. Arakaki T, Mahon S, Charpier S, Leblois A, Hansel D. The Role of Striatal Feedforward Inhibition in the Maintenance of Absence Seizures. J Neurosci. 2016;36(37):9618–9632.

69. Chen M, Guo D, Wang T, Jing W, Xia Y, Xu P, et al. Bidirectional Control of Absence Seizures by the Basal Ganglia: A Computational Evidence. PLoS Comput Biol. 2014;10(3):1–17.

70. Chen M, Guo D, Li M, Ma T, Wu S, Ma J, et al. Critical Roles of the Direct GABAergic Pallido-cortical Pathway in Controlling Absence Seizures. PLoS Comput Biol. 2015;11(10):1–23.

